# Deep Learning Meets Sleep Medicine: A Proof-of-Concept Clustering of Minute-Resolution CPAP Telemetry

**DOI:** 10.1101/2025.07.03.663061

**Authors:** Latherial Calbert, Matthew T. Scharf, Ioannis P. Androulakis

## Abstract

This proof-of-concept study demonstrates that minute-resolution telemetry from continuous positive airway pressure (CPAP) devices can be effectively repurposed for large-scale chronotype and adherence phenotyping. We collated 30 consecutive nights from *n*=200 de-identified ResMed patients into 30 *×* 1440 colour images, embedded each image with a frozen ResNet-50 convolutional neural network, and clustered the embeddings with *k* -means. Six distinct phenotypes emerged, capturing both sleep timing (early birds, typical sleepers, night owls) and adherence patterns (high, medium-high, inconsistent, fragmented, and non-adherent). The approach leverages routinely collected clinical data without the need for additional sensors, promising significant benefits for personalized sleep medicine. External validation against actigraphy and questionnaire-based chronotype measures is planned to further strengthen these findings.

## 1 Introduction

Obstructive sleep apnea (OSA) affects an estimated 9–38% of adults worldwide [Senaratna et al., 2017]. Continuous positive airway pressure (CPAP) therapy remains the gold standard but suffers from poor adherence [Weaver and Grunstein, 2010]. Recent machine learning approaches have shown promise in predicting CPAP adherence, with studies achieving F1 scores up to 0.864 [Ikotun et al., 2022] and demonstrating that personalized ML-driven interventions can substantially increase adherence [Lacroix et al., 2023]. However, these approaches typically focus on predicting adherence outcomes rather than understanding the underlying phenotypes that drive usage patterns. Because modern devices record mask status every minute for insurance compliance, they inadvertently capture rich temporal information about each user’s sleep timing. Unlocking these data could furnish clinicians with chronotype insights at scale and with zero patient burden, revolutionizing personalized approaches to sleep medicine.

Machine learning offers a powerful means to uncover structure in such high-dimensional, temporally rich datasets. In particular, unsupervised learning techniques such as clustering can reveal naturally occurring usage patterns that may be obscured by conventional summary statistics like average nightly use. These patterns are of clinical interest: they may reflect not only treatment engagement but also underlying circadian phenotypes—critical for tailoring interventions and predicting long-term adherence trajectories. Indeed, chronotype—the individual preference for sleep-wake timing—has been increasingly recognized as an important factor in health outcomes, with evening chronotypes showing higher risk for various health conditions [Zou et al., 2022].

Traditional approaches to analyzing CPAP adherence data have relied on extracting numerical features from usage logs, such as mean hours per night or percentage of nights with adequate use. However, these approaches may miss complex temporal patterns that are readily apparent in visual representations of the data. The human visual system excels at pattern recognition, and by transforming CPAP usage data into images, we can leverage powerful computer vision techniques that have been pre-trained to extract meaningful features from visual inputs. This image-based approach offers several advantages: (1) it preserves the full temporal structure of the data without requiring assumptions about which features matter; (2) it requires minimal preprocessing and avoids the need for manual feature engineering; (3) it can capture subtle patterns that might be difficult to encode as explicit features, such as gradual shifts in sleep timing or weekend-weekday differences; and (4) it enables the use of pre-trained deep learning models that have already learned to extract hierarchical visual features from millions of images.

The visual representation of temporal data as images has precedent in other domains. For example, recurrence plots have been used to visualize dynamical systems [Eckmann et al., 1987], and spectrograms convert audio signals into visual representations for analysis. By rendering each patient’s 30-day CPAP usage as a heat-map image, we create a visual “fingerprint” that captures their unique usage pattern in a format amenable to computer vision analysis. This approach transforms the problem from time-series analysis to image classification, allowing us to leverage the remarkable advances in deep learning for computer vision.

In a novel application of machine learning to CPAP adherence data, Scharf and Androulakis [Scharf and Androulakis, 2025] developed a methodology to characterize and cluster diurnal patterns of CPAP use among patients with obstructive sleep apnea (OSA). By transforming 30-day adherence data into a standardized 24-hour usage profile and approximating individual patterns with four parameters—onset time, offset time, and usage consistency within and outside that window—the study identified six distinct adherence-timing clusters in a cohort of 200 patients. Their approach distinguishes between normal and delayed timing patterns and stratifies patients by adherence level, highlighting the potential of routinely collected CPAP data as a proxy for diurnal rhythmicity. This work represents a significant step toward understanding the role of treatment timing in modulating CPAP efficacy and underscores the need to incorporate temporal adherence patterns into clinical assessment. Building on this work, the present study advances the analysis by incorporating higher-resolution temporal data to capture finer-grained fluctuations and dynamic patterns of CPAP use.

Deep learning, specifically convolutional neural networks (CNNs), has demonstrated exceptional ability to extract meaningful features from complex input domains such as time-series data, especially when re-rendered as images. By transforming CPAP usage into visual representations and leveraging pre-trained CNNs, we can repurpose widely available, passive adherence data into actionable chronobiological insights—without requiring additional sensors, patient surveys, or behavioral diaries. While artificial intelligence has been increasingly applied to OSA treatment optimization [Brennan and Kirby, 2023], our approach uniquely combines visual pattern recognition with unsupervised phenotype discovery.

In this proof-of-concept study, we demonstrate that minute-resolution CPAP telemetry, analyzed with standard deep learning and clustering tools, can identify distinct and interpretable adherence–chronotype phenotypes. This approach could enable precision sleep medicine at scale, offering a low-burden, high-impact path to improving patient stratification and care.

## 2 Methods

### 2.1 Data Source

A retrospective cohort study was conducted on patients treated at the Comprehensive Sleep Disorders Center at Rutgers Robert Wood Johnson Medical School between 2013 and 2023. In accordance with insurance criteria, patients with an AHI greater than or equal to 15, or AHI greater than or equal to 5 with a qualifying comorbidity, were offered CPAP therapy. De-identified CPAP adherence data for 200 patients were obtained from the ResMed AirView^®^ platform, which records CPAP use within 24-hour periods starting at midnight. To avoid variability during the initial acclimation phase, data from days 61–90 of therapy were analyzed. Patients lacking data for this period or with more than 12 CPAP on/off cycles per day were excluded due to data unreliability. A 30-day time horizon was selected arbitrarily but did not affect the analytical method. Data were collected sequentially from 200 records. The study was approved by the Rutgers IRB (Pro2023001290), and all ethical standards were upheld.

### 2.2 Image Construction

For each patient we stacked 30 consecutive nights into a 30 *×* 1440 matrix and rendered it as a pseudo-colour PNG. Rows represent successive calendar days, while columns sweep chronologically across the 24-hour period: the leftmost column marks midnight and the rightmost column 23:59. In these visual representations, white denotes nighttime periods, yellow denotes daytime periods, and red horizontal bars indicate when the CPAP mask was in use. This color scheme was chosen to provide clear visual contrast and facilitate pattern recognition. Colour scaling was standardized across all patients. Original heat-map images are 875*×* 656 pixels before processing and are isotropically resized to 224 *×* 224 pixels for compatibility with ResNet-50.

### 2.3 Feature Extraction and Preprocessing

A frozen ResNet-50 pre-trained on ImageNet [He et al., 2016] generates a 2,048-dimensional embedding from the global-average-pooling layer. We chose this pre-trained CNN approach because it leverages the powerful feature extraction capabilities developed through training on millions of images, eliminating the need for specialized algorithm development. The network’s internal representations have been shown to capture complex visual patterns that transfer well to diverse domains. No post-hoc min–max scaling is applied beyond the network’s internal normalization; no colour jittering or augmentation is used.

#### 2.3.1 Outlier Detection

Prior to clustering analysis, we applied Isolation Forest [Liu et al., 2008] for outlier detection to ensure robust clustering results. Isolation Forest is particularly well-suited for high-dimensional data as it isolates anomalies by randomly selecting features and split values, requiring fewer partitions to isolate outliers compared to normal points [Liu et al., 2012]. This preprocessing step is crucial because clustering algorithms, particularly k-means, are sensitive to outliers that can distort cluster centers and lead to suboptimal partitioning [Chandola et al., 2009]. In our analysis, Isolation Forest identified and removed 5 outliers from the dataset. However, for clinical completeness and to maintain our cohort size of 200 patients, these outliers were retained in the final visualization and clustering analysis, ensuring that our phenotypes represent the full spectrum of CPAP usage patterns encountered in clinical practice.

#### 2.3.2 Dimensionality Reduction

Following outlier removal, we applied Principal Component Analysis (PCA) to reduce the 2,048-dimensional ResNet embeddings while retaining 95% of the variance. PCA serves multiple critical purposes in our pipeline: (1) it reduces computational complexity for subsequent clustering, making the analysis more efficient [Jolliffe, 2002]; (2) it mitigates the curse of dimensionality that can affect distance-based clustering algorithms in high-dimensional spaces [Beyer et al., 1999]; and (3) it acts as a denoising step by concentrating signal in the first principal components while relegating noise to components with lower variance [Vidal et al., 2016]. This dimensionality reduction is particularly important for k-means clustering, which relies on Euclidean distances that can become less meaningful in very high-dimensional spaces.

The combination of outlier removal and PCA creates a robust preprocessing pipeline that ensures our clustering analysis focuses on the most informative patterns in the data while being computationally efficient and resistant to noise and anomalies.

### 2.4 Dimensionality Reduction

Two-dimensional t-SNE via scikit-learn’s default parameters (perplexity = 30, learning rate = 200, n_iter = 1000, random_state = 42) visualizes the embedding space. We selected t-SNE for dimensionality reduction because it excels at preserving local structure while revealing global patterns in high-dimensional data. The t-SNE projection proved particularly valuable for revealing the natural structure within our data, providing compelling visual evidence of distinct usage patterns that would be difficult to discern in the original 2,048-dimensional space.

### 2.5 Clustering and *k* Selection

To determine the optimal number of clusters, we employed a comprehensive validation approach using four complementary metrics selected based on recent clustering validation research [Arbelaitz et al., 2013, Halkidi and Vazirgiannis, 2001]:

#### 1. C-Index (Concordance Index)

A measure that compares the sum of distances between points within clusters to the minimum and maximum possible sums of distances in the dataset. Specifically, it calculates (*S − S*_*min*_)*/*(*S*_*max*_ *− S*_*min*_) where *S* is the sum of within-cluster distances, *S*_*min*_ is the sum of the smallest distances, and *S*_*max*_ is the sum of the largest distances. Values range from 0 to 1, with lower values indicating better clustering [Hubert and Arabie, 1985].

#### 2. Silhouette Coefficient

Measures how similar each point is to its own cluster compared to neighboring clusters. Values range from -1 to 1, where higher values indicate well-separated and compact clusters [Rousseeuw, 1987].

#### 3. Davies-Bouldin Index

Evaluates the average similarity between each cluster and its most similar cluster, where similarity accounts for both within-cluster scatter and between-cluster separation. Lower values indicate better clustering [Davies and Bouldin, 1979].

#### 4. S Dbw (Scatter and Density between clusters)

A density-based validation index that considers both intra-cluster compactness and inter-cluster separation using density measures. Lower values indicate better clustering quality [Halkidi and Vazirgiannis, 2001].

We selected these four metrics to provide a comprehensive evaluation covering different aspects of clustering quality: compactness (C-Index), separation (Silhouette), similarity (Davies-Bouldin), and density-based structure (S Dbw). This multi-metric approach ensures robust validation beyond any single criterion [Arbelaitz et al., 2013].

For the clustering range, we tested *k* values from 3 to 8 clusters. This range was chosen based on established practices in clustering validation literature, which suggest that testing extreme values (very few or very many clusters) often yields suboptimal results [Thorndike, 1953, Yuan and Yang, 2019]. The lower bound of 3 ensures meaningful segmentation beyond simple binary divisions, while the upper bound of 8 prevents over-fragmentation that could obscure interpretable patterns in clinical data [Ketchen and Shook, 1996].

K-means remains one of the most widely used clustering algorithms due to its computational efficiency and interpretability, particularly suitable for large-scale applications [Ikotun et al., 2023]. To address the sensitivity of k-means to initialization, we ran the algorithm 10 times with different random seeds for each value of *k*.

For each of the four metrics, we calculated the mean score across the 10 runs and identified the *k* value that optimized that metric (minimum for C-Index, Davies-Bouldin, and S Dbw; maximum for Silhouette). While the individual metrics suggested different optimal values—C-Index indicated *k* = 5, Silhouette suggested *k* = 3, and both Davies-Bouldin and S Dbw pointed to *k* = 8—the average of these four values yielded our final recommendation of 6 clusters: (5 + 3 + 8 + 8)*/*4 = 6.

The t-SNE visualization independently supported this finding through visual inspection, with six distinct islands clearly visible in the embedding space (Fig. 2). This dual validation—both quantitative metrics and qualitative visualization—strengthens our confidence in the identified sleep-timing and adherence phenotypes.

## 3 Results

### 3.1 Optimal Cluster Count

Figure 1 summarises model-selection curves across our four evaluation metrics for cluster counts ranging from 3 to 8. We employed a rigorous validation approach: for each value of *k*, we ran k-means clustering 10 times with different random initializations to account for the algorithm’s sensitivity to starting conditions.

**Figure 1:**
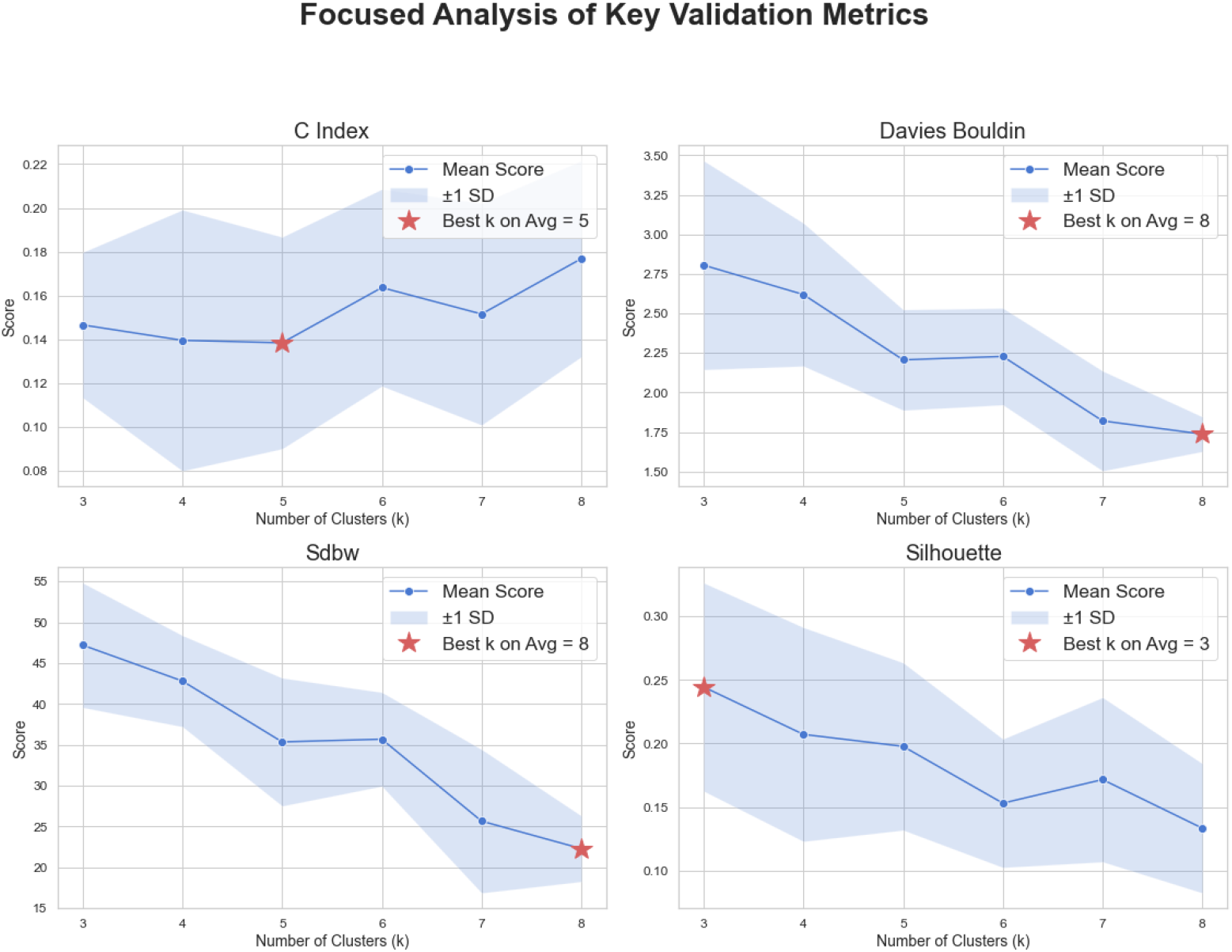
Focused analysis of key validation metrics for *k* = 3 … 8 displayed in a 2×2 matrix. Each panel shows the mean score (solid line with markers) and ±1 standard deviation (shaded area) across 10 independent k-means runs with different initializations. Red stars indicate the optimal *k* value for each metric. (Top left) C-Index shows optimal clustering at *k* = 5. (Top right) Davies-Bouldin index reaches its minimum at *k* = 8. (Bottom left) S Dbw metric indicates optimal clustering at *k* = 8. (Bottom right) Silhouette coefficient peaks at *k* = 3. While individual metrics suggest different optimal values, their average (5 + 3 + 8 + 8)*/*4 = 6 provides a balanced recommendation of six clusters that captures the essential structure in the CPAP usage data.

For each metric across the 10 runs, we calculated the mean and standard deviation of scores. The optimal *k* for each metric was determined based on these averaged scores: the C-Index reached its minimum at *k* = 5, the Silhouette coefficient showed its peak at *k* = 3, while both the Davies-Bouldin index and S Dbw metric indicated *k* = 8 as optimal. Despite this variation, these values cluster around the mid-range of our tested values.

To obtain our final recommendation, we calculated the average of these four optimal *k* values: (5 + 3 + 8 + 8)*/*4 = 6. This averaging approach balances the different perspectives each metric provides—C-Index and Silhouette emphasizing geometric separation, while Davies-Bouldin and S Dbw focusing on density-based clustering quality. The resulting *k* = 6 represents a robust compromise that captures the essential structure in the data while avoiding over-fragmentation. The shaded regions in Figure 1 show ±1 standard deviation across the 10 initialization runs, demonstrating relatively consistent performance despite the stochastic nature of k-means initialization.

### 3.2 t-SNE Visualization

Figure 2 shows the 2-D t-SNE embedding of all 200 patients, where six distinct islands are readily apparent. Each thumbnail within the visualization represents one patient’s 30-day CPAP usage pattern. The clear spatial separation between clusters in the embedding space provides compelling visual evidence that our image-based approach captures meaningful differences in CPAP usage patterns. The clustering assignments (indicated by colored borders) align remarkably well with the natural groupings visible in the t-SNE projection, validating our choice of six clusters.

**Figure 2:**
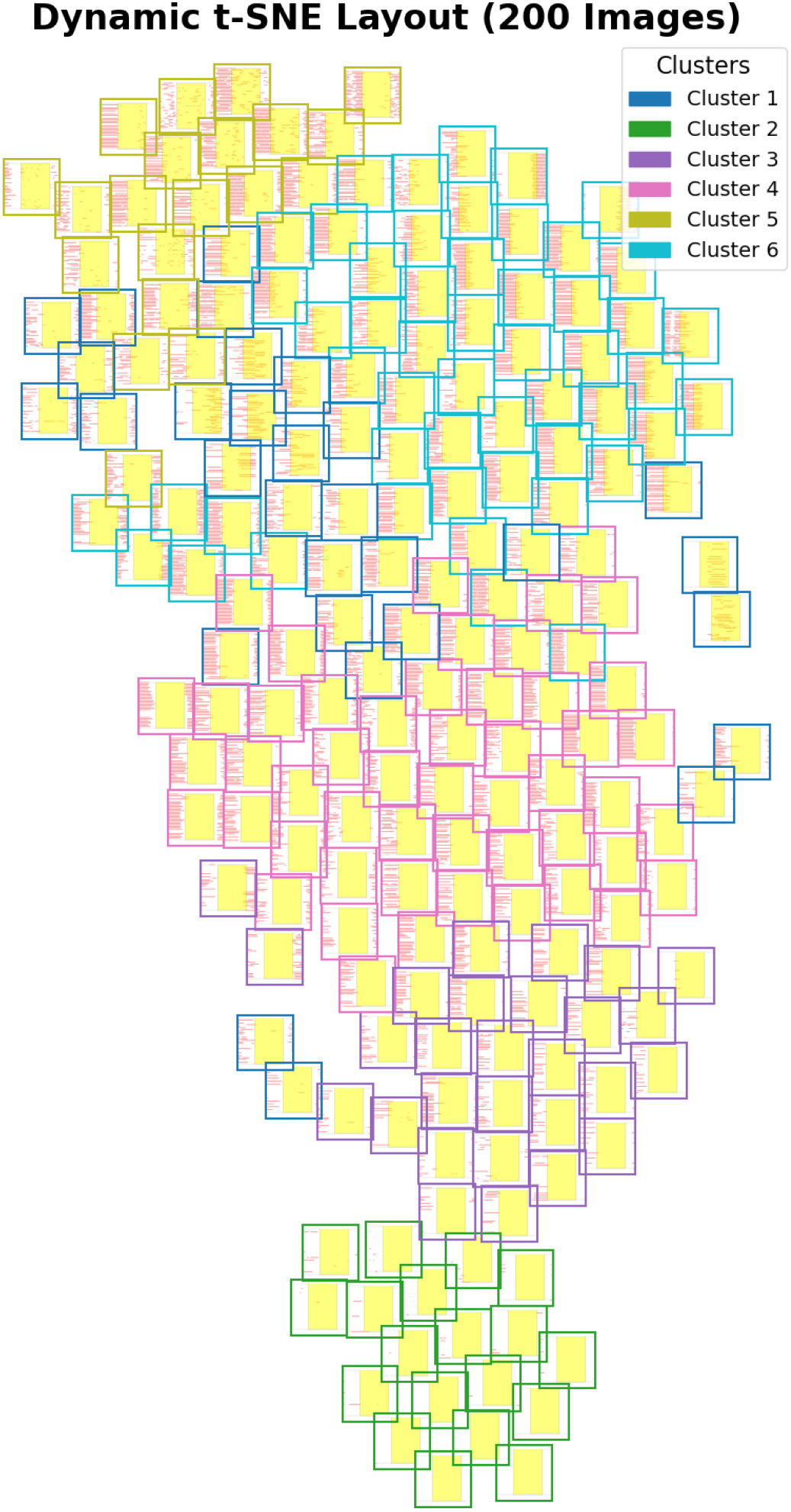
t-SNE projection of ResNet embeddings reveals six distinct clusters among 200 CPAP users. The visualization displays all 200 patient heat-maps as thumbnails (note: 5 outliers were removed by Isolation Forest during preprocessing, but the final analysis includes the complete set of 200 patients for clinical relevance). Colored borders encircle each cluster (Clusters 1-6), demonstrating the natural groupings discovered by our analysis. Within each thumbnail heat-map, rows represent consecutive days (top to bottom) and columns represent time of day (midnight to midnight, left to right). The clear spatial separation between the six encircled groups validates our clustering approach and demonstrates that visual patterns of CPAP usage encode meaningful phenotypic information. The large separation distances between clusters indicate that these represent genuinely distinct usage patterns rather than arbitrary divisions of a continuum.

### 3.3 Cluster Characterization

Representative 30-day heat-maps for each cluster appear in Fig. 3. Through visual inspection of all cluster members (see Appendix for complete results), we identified distinct combinations of sleep timing and adherence quality. Each horizontal red band within a heat-map corresponds to one night of CPAP use, with time of day running left-to-right from midnight to midnight. White regions indicate nighttime hours, yellow indicates daytime, and red bars show periods of active CPAP use.

**Figure 3:**
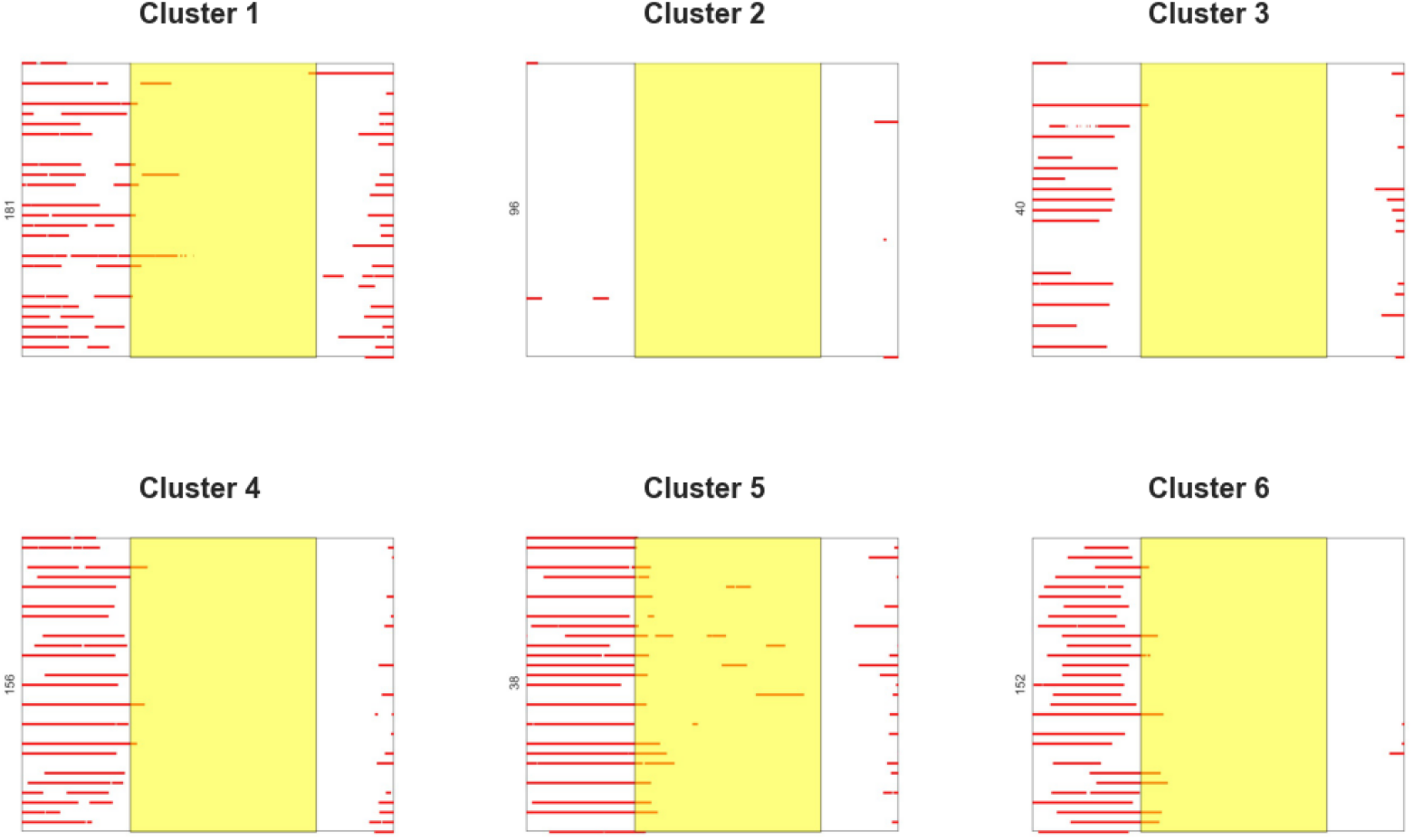
Representative CPAP usage patterns showing one exemplar patient from each of the six identified clusters, arranged in a 2×3 grid. Each panel displays a single patient’s 30-day usage pattern that best represents their cluster’s characteristics (selected as the patient closest to the cluster centroid). Color coding: yellow = daytime hours (6 AM-6 PM), white = nighttime hours (6 PM-6 AM), red = CPAP mask on. Time progresses from midnight to midnight (left to right), with each horizontal stripe representing one calendar day. Visual inspection reveals distinct phenotypes: Cluster 1 shows consistent early evening adherence; Cluster 2 shows minimal usage; Cluster 3 displays typical sleep timing with moderate adherence; Cluster 4 exhibits consistent usage with standard timing; Cluster 5 shows severe fragmentation; Cluster 6 demonstrates delayed sleep onset with variable adherence. These representative examples guide readers to the complete cluster membership shown in the Appendix.

## 4 Discussion

This proof-of-concept study demonstrates the power of combining computer vision techniques with unsupervised learning to extract clinically meaningful phenotypes from routine CPAP telemetry. CPAP telemetry, defined as the remote collection, transmission, and analysis of data from CPAP devices, has the potential to emerge as a critical tool in modern sleep medicine, enabling continuous access to detailed usage metrics such as nightly adherence, mask leak rates, and residual respiratory events. By transforming this temporal usage data into visual representations and leveraging pre-trained deep learning models, we identified six distinct clusters that capture both circadian timing and adherence quality without requiring any manual feature engineering. This approach not only illustrates the potential of automated, data-driven phenotyping, but also highlights the broader value of CPAP telemetry in improving clinical oversight, facilitating early intervention, and generating actionable insights at both individual and population levels.

The decision to analyze CPAP data as images rather than traditional time-series represents a paradigm shift with several key advantages. Traditional approaches require researchers to predefine relevant features (e.g., mean usage hours, standard deviation of bed-time), potentially missing complex patterns that don’t fit predetermined metrics. In contrast, our visual approach preserves all temporal information in an intuitive format that captures patterns a human expert would recognize—but at scale and with mathematical rigor. The pre-trained ResNet-50 model brings sophisticated pattern recognition capabilities developed from millions of natural images, detecting subtle visual features that might escape manual analysis. This transfer learning approach means we benefit from years of computer vision research without developing CPAP-specific algorithms from scratch.

Our multi-metric validation approach, combining C-Index, Silhouette coefficient, Davies-Bouldin index, and S Dbw across multiple initialization runs, provided robust quantitative evidence for our clustering solution. While individual metrics suggested different optimal cluster numbers (ranging from 3 to 8), their average converged on six clusters, representing a balanced solution that avoids both under-segmentation and over-fragmentation. This finding was independently supported through visual inspection of the t-SNE projection, which revealed six clearly separated islands in the embedding space.

Through systematic visual inspection of all cluster members (see Appendix), we characterized the six phenotypes as follows:

### 1. Early birds with excellent adherence

(Cluster 1, n=30): These patients demonstrate consistent early evening sleep onset (typically 9:00-10:00 PM) with sustained usage for 7-9 hours nightly. The visual pattern shows uniform red bands starting early in the evening and continuing until 5:00-6:00 AM. These patients likely have traditional work schedules and prioritize sleep hygiene. Their high adherence (*>* 95% of nights with *>* 6 hours use) suggests excellent therapy acceptance and minimal side effects. This phenotype represents the ideal treatment outcome.

### 2. Non-adherent patients

(Cluster 2, n=19): This cluster shows minimal to no CPAP usage (*<* 10% of nights with meaningful use). The heat-maps are predominantly white and yellow with sporadic, brief red marks lasting *<* 2 hours when present. These patients have essentially abandoned therapy within the monitoring period. Visual inspection reveals some attempted use in early days, suggesting initial effort followed by discontinuation. This phenotype requires immediate intervention to identify barriers (mask discomfort, claustrophobia, perceived lack of benefit) and consider alternative therapies.

### 3. Typical sleepers with moderate adherence

(Cluster 3, n=28): Patients display conventional sleep timing (11:00 PM-12:00 AM onset, 6:00-7:00 AM offset) with 70-85% nightly usage. Visual inspection reveals regular patterns with occasional gaps, particularly on weekends. The missing nights appear random rather than systematic, suggesting forgetfulness or situational factors rather than fundamental issues with therapy. These patients would benefit from behavioral interventions and reminder systems to achieve optimal adherence.

### 4. Consistent users with standard timing

Cluster 4, n=51): The largest cluster shows highly consistent CPAP usage (*>* 90% of nights) with conventional sleep timing (10:30 PM-6:30 AM). The visual pattern displays remarkably uniform red bands with minimal variation, indicating ingrained habit formation. Weekend-weekday differences are negligible, suggesting lifestyle stability. This represents successful long-term adherence achieved through routine establishment.

### 5. Fragmented sleep patterns

(Cluster 5, n=20): These heat-maps show multiple on/off cycles within nights (3-8 segments), appearing as broken red segments rather than continuous bands. Average total usage may reach 4-6 hours but distributed across the night. This pattern suggests either: (a) severe sleep fragmentation from untreated comorbidities (PLM, central apnea); (b) mask/pressure tolerance issues causing repeated awakenings; or (c) nocturia or other medical conditions. The inconsistent timing of fragments indicates underlying sleep architecture disruption requiring comprehensive evaluation.

### 6. Night owls with variable adherence

(Cluster 6, n=52): This cluster exhibits consistently late sleep onset (1:00-3:00 AM) with wake times extending to 9:00-11:00 AM. Usage quality varies considerably: approximately 40% maintain good adherence (*>* 6 hours) within their delayed window, while 60% show irregular patterns with 3-5 hour usage. The heat-maps reveal two sub-patterns: stable night owls who consistently use CPAP during their preferred hours, and unstable night owls with erratic usage. This suggests that chronotype alone doesn’t determine adherence success—schedule consistency matters more than timing.

These phenotypes reveal critical insights for personalized medicine. Visual inspection of the heat-maps shows a clear distinction between Clusters 3 and 4, demonstrating that moderate versus excellent adherence isn’t simply about hours of use but pattern consistency. Both groups use CPAP most nights, but Cluster 4’s unwavering routine (visible as perfectly aligned red bands) contrasts with Cluster 3’s occasional lapses (gaps in the heat-map). This visual analysis suggests that interventions should focus on habit formation rather than just education about benefits.

The fragmented pattern of Cluster 5 is visually distinct from all others—while poor adherence in Cluster 2 shows absence of use, Cluster 5 shows presence of effort undermined by repeated interruptions. This distinction, easily visible in heat-maps but potentially missed in summary statistics, has important clinical implications: Cluster 2 needs motivation and barrier removal, while Cluster 5 needs medical optimization.

Visual temporal analysis reveals that successful users (Clusters 1, 4, and stable members of 6) typically initiate CPAP within 15 minutes of bedtime and maintain usage until natural awakening, while struggling users show delays of 30-90 minutes between bedtime and mask application, suggesting bedtime procrastination or anxiety about therapy.

The image-based approach offers several advantages over traditional numerical feature extraction. First, it preserves the complete temporal structure of the data without requiring assumptions about which features matter. Second, it leverages the sophisticated pattern recognition capabilities of pre-trained CNNs, which have learned to extract hierarchical visual features from millions of images. Third, it provides intuitive visualizations that clinicians can immediately interpret, bridging the gap between complex machine learning and clinical practice.

Our findings align with recent work showing chronotype’s impact on sleep parameters [Colelli et al., 2023], but extend beyond simple timing classifications. The visual patterns reveal that adherence quality varies independently of sleep timing—we observe both successful and struggling patients across different chronotypes. This suggests that personalized interventions should consider both when patients prefer to sleep and how consistently they can maintain CPAP usage within their preferred window.

The clinical implications are substantial. For Clusters 1 and 4 (consistent users), the focus should be on maintenance and positive reinforcement. Cluster 3 patients might benefit from targeted adherence coaching to improve consistency. Cluster 6 (night owls) may need schedule accommodations rather than attempts to shift their sleep timing. Cluster 5 requires investigation for mask fit, pressure settings, or comorbid conditions. Cluster 2 needs comprehensive re-evaluation of their treatment approach, possibly including alternative therapies.

Our visual pattern recognition approach complements recent ML efforts in CPAP adherence prediction [Ikotun et al., 2022, Lacroix et al., 2023] by focusing on phenotype discovery rather than outcome prediction. This methodology could be integrated with existing clinical decision support systems to provide real-time phenotype identification and guide personalized interventions.

#### Limitations

Several limitations should be acknowledged: (1) Demographic and clinical outcome data were unavailable for correlation with clusters; (2) External validation against established chronotype measures (e.g., Morningness-Eveningness Questionnaire) is still needed; (3) The visual inspection approach, while clinically intuitive, could benefit from quantitative feature extraction to complement the qualitative characterization; (4) The sample size (n=200), while substantial for proof-of-concept, should be expanded to ensure generalizability.

Future work will validate clusters against actigraphy data, questionnaire-based chronotype assessments, and long-term clinical outcomes. We also plan to extract quantitative metrics (mean onset time, usage duration, fragmentation index) for each cluster to complement the visual characterization.

## 5 Conclusion

This study demonstrates that minute-resolution CPAP telemetry, when transformed into color-coded heat-maps and analyzed using pre-trained deep learning models, reveals six clinically meaningful phenotypes. The image-based approach leverages powerful computer vision techniques while maintaining clinical interpretability. By identifying distinct combinations of sleep timing and adherence quality, this method enables personalized interventions tailored to each patient’s specific usage pattern.

Our findings demonstrate high confidence in the identified phenotypes through multiple validation approaches. The four diverse clustering metrics (C-Index, Silhouette, Davies-Bouldin, and S Dbw) suggested different optimal cluster numbers (5, 3, 8, and 8 respectively), but their average converged on six clusters—a balanced solution that captures the essential structure without over-fragmentation. Each metric evaluates different aspects of clustering quality, and their collective assessment provides more robust validation than any single criterion. This statistical validation is reinforced by the clear visual separation observed in t-SNE projections and the clinical coherence of the identified patterns through systematic visual inspection of all heat-maps. The phenotypes align with known sleep medicine concepts while revealing nuanced distinctions—such as fragmented versus absent usage—that traditional metrics might miss.

The approach’s strength lies in its ability to transform routine clinical data into actionable insights without additional patient burden. Every CPAP machine already collects minute-by-minute usage data; our method simply views this data through a new lens. The visual representations provide intuitive understanding for clinicians while the deep learning analysis ensures objective, reproducible phenotyping.

Future work should validate these phenotypes against clinical outcomes, including cardiovascular events, daytime sleepiness scores, and quality of life measures. Prospective studies could test whether phenotype-tailored interventions (e.g., chronotherapy for night owls, sleep consolidation therapy for fragmented users) improve outcomes compared to standard care. Integration with electronic health records could enable real-time phenotype identification and trigger appropriate interventions. Additionally, extending this approach to other sleep disorders and home monitoring devices could establish visual pattern recognition as a general paradigm in sleep medicine.

As sleep medicine embraces precision approaches, visual pattern recognition of routine device data offers a powerful tool for improving patient outcomes. By revealing the hidden structure in CPAP usage patterns, this method transforms adherence monitoring from a simple compliance metric to a window into patients’ sleep-wake physiology and behavior. The approach requires no additional sensors or patient effort, making it immediately scalable to the millions of CPAP users worldwide.

## Data and Code Availability

All analysis scripts and an example de-identified dataset are available at https://github.com/latherialcalbert/cpap-adherence-clustering-resnet50-kmeans.

## Acknowledgements

LC acknowledges support from NSF (EEC-23-49225) REU Site - Cellular Bioengineering: From Biomaterials to Stem Cells.

## Appendix: Complete Cluster Membership

The following figures present all 200 patients organized by their cluster membership. Each heat-map represents one patient’s 30-day CPAP usage pattern during days 61-90 of therapy. Within each heat-map:

- Rows represent consecutive days (top to bottom)
- Columns represent time of day progressing from midnight to midnight (left to right)
- Yellow indicates daytime hours (6:00 AM - 6:00 PM)
- White indicates nighttime hours (6:00 PM - 6:00 AM)
- Red bars indicate periods of active CPAP use

The figures are organized to avoid overcrowding, with no more than 30-35 patients displayed per page. This comprehensive visualization allows readers to appreciate both the within-cluster consistency and between-cluster differences that define our six phenotypes.

**Cluster 1:**
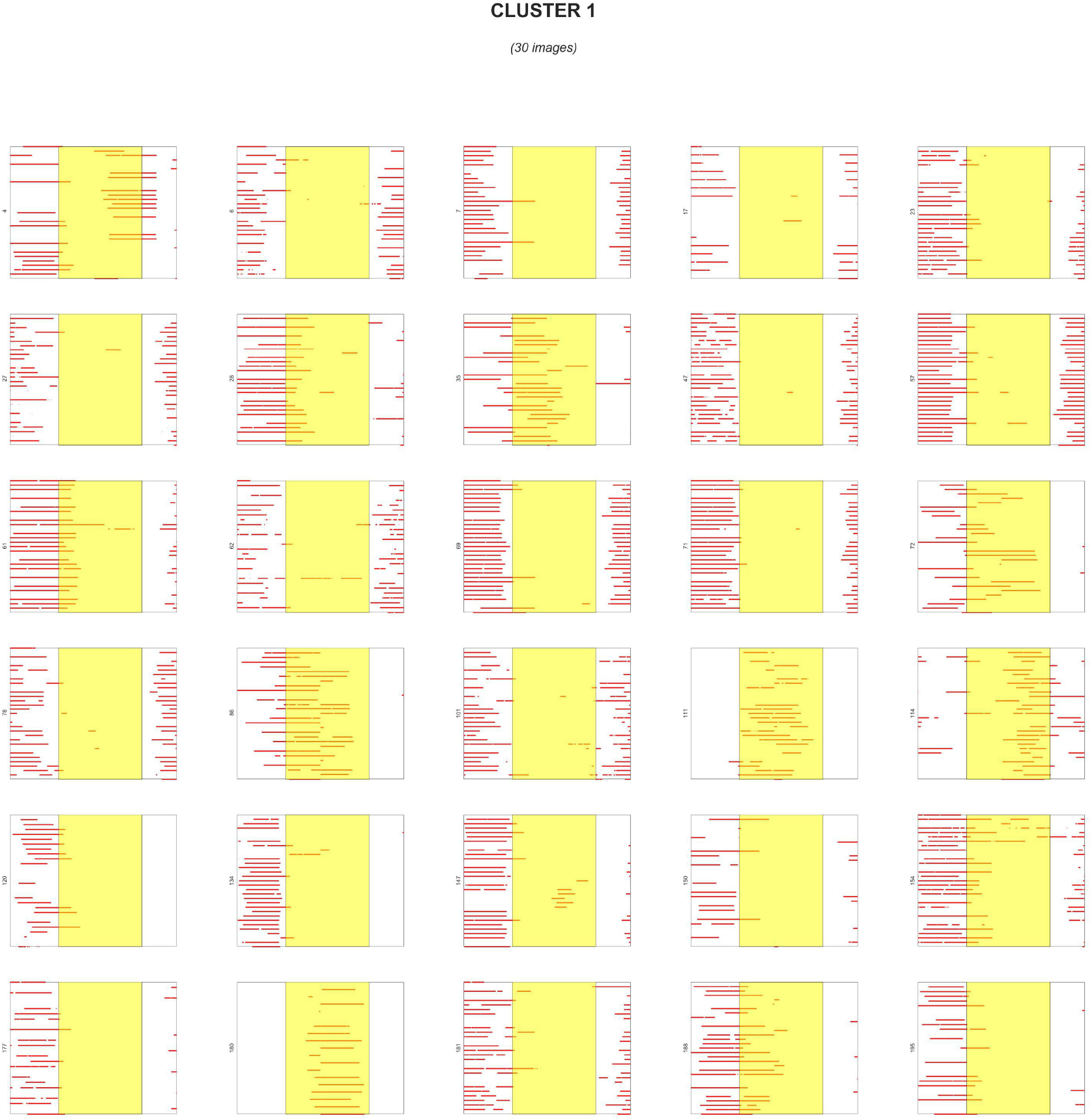
Early birds with excellent adherence (n=30). All patients in this cluster demonstrate consistent early evening sleep onset (9:00-10:00 PM) with sustained usage throughout the night. Note the remarkable uniformity of red bands across all patients, indicating stable sleep schedules and excellent therapy adherence.

**Cluster 2:**
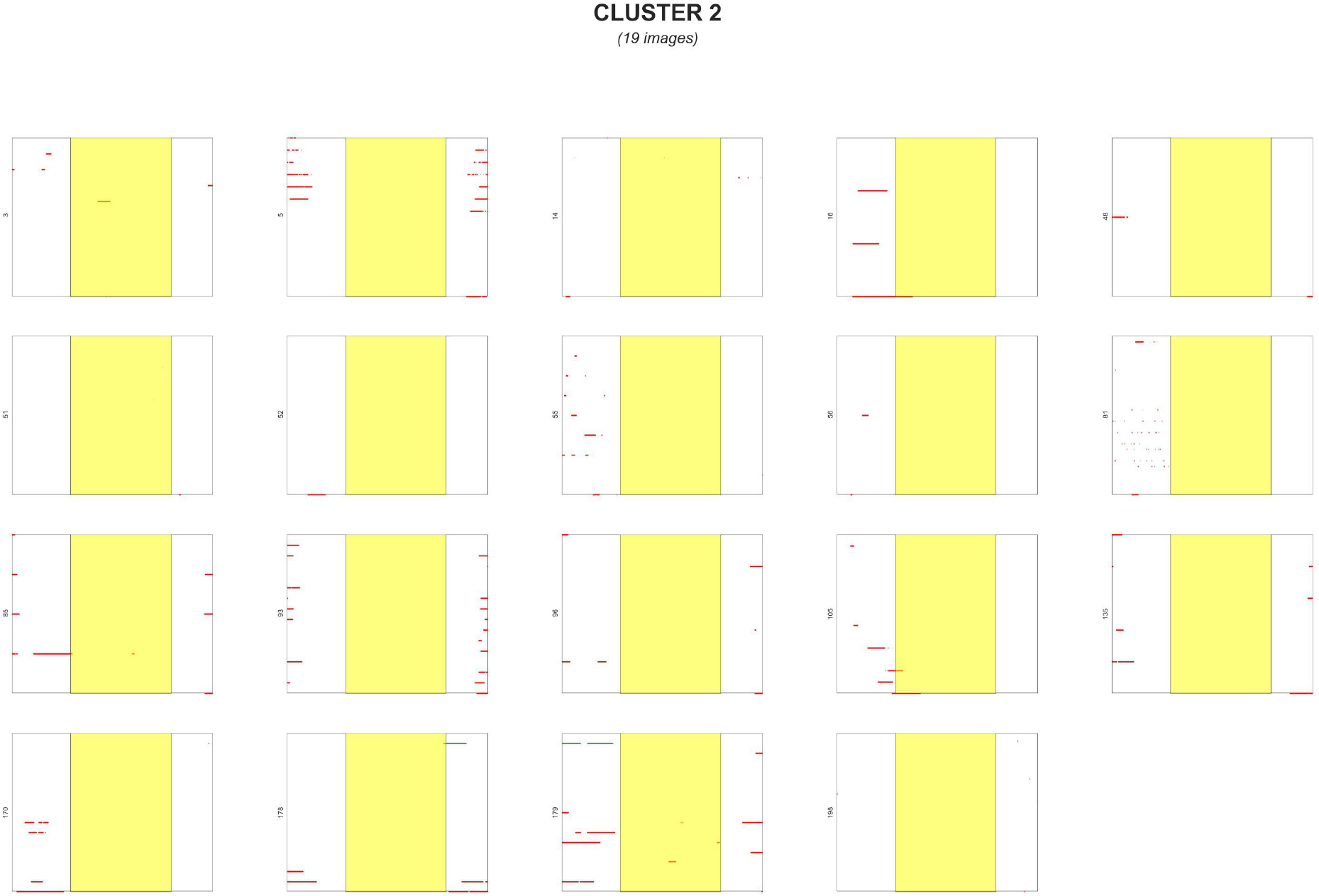
Non-adherent patients (n=19). These patients show minimal to no CPAP usage, requiring immediate clinical intervention.

**Cluster 3:**
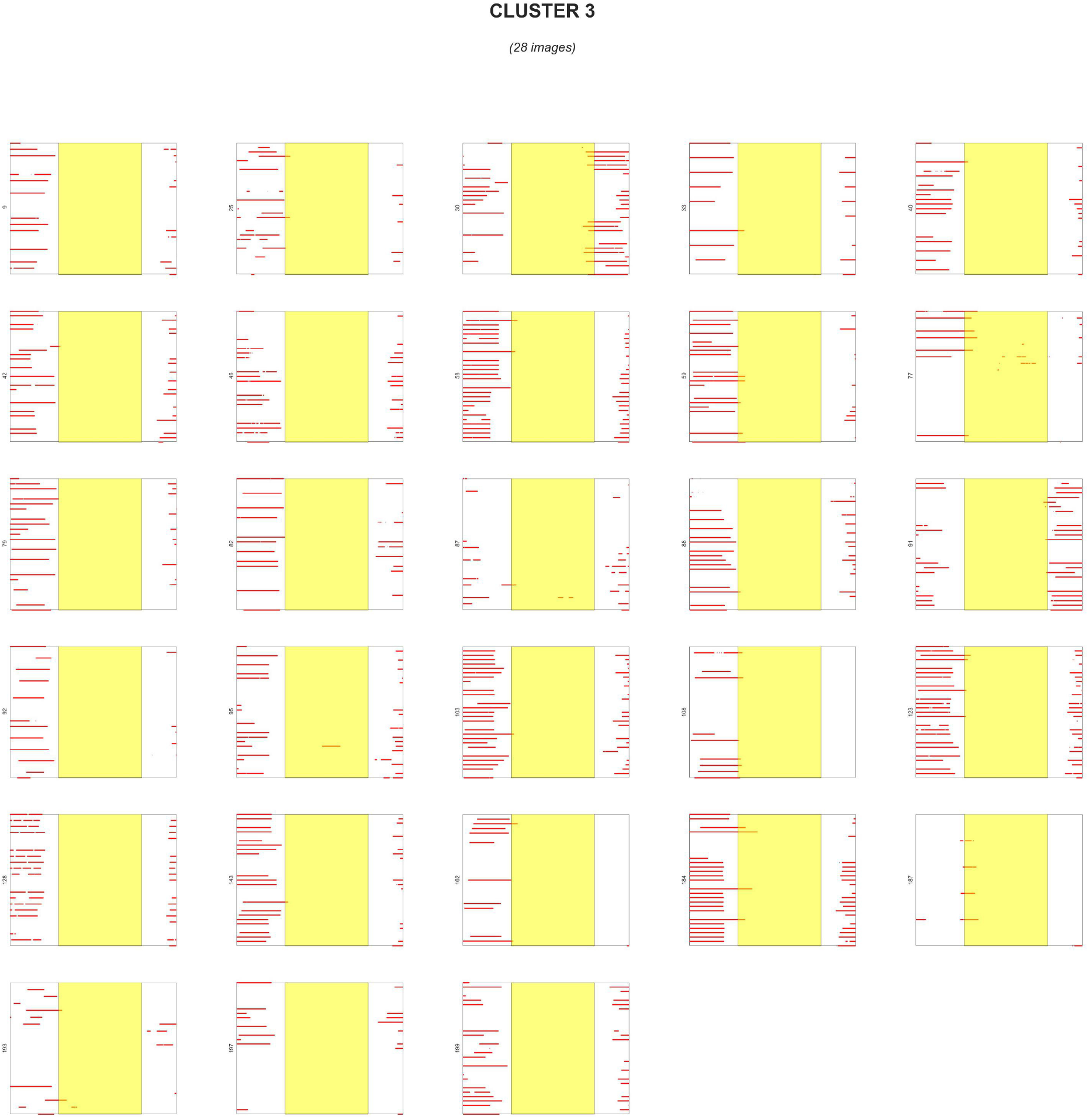
Typical sleepers with moderate adherence (n=28). Patients display intermediate sleep timing with good but improvable adherence patterns.

**Cluster 4:**
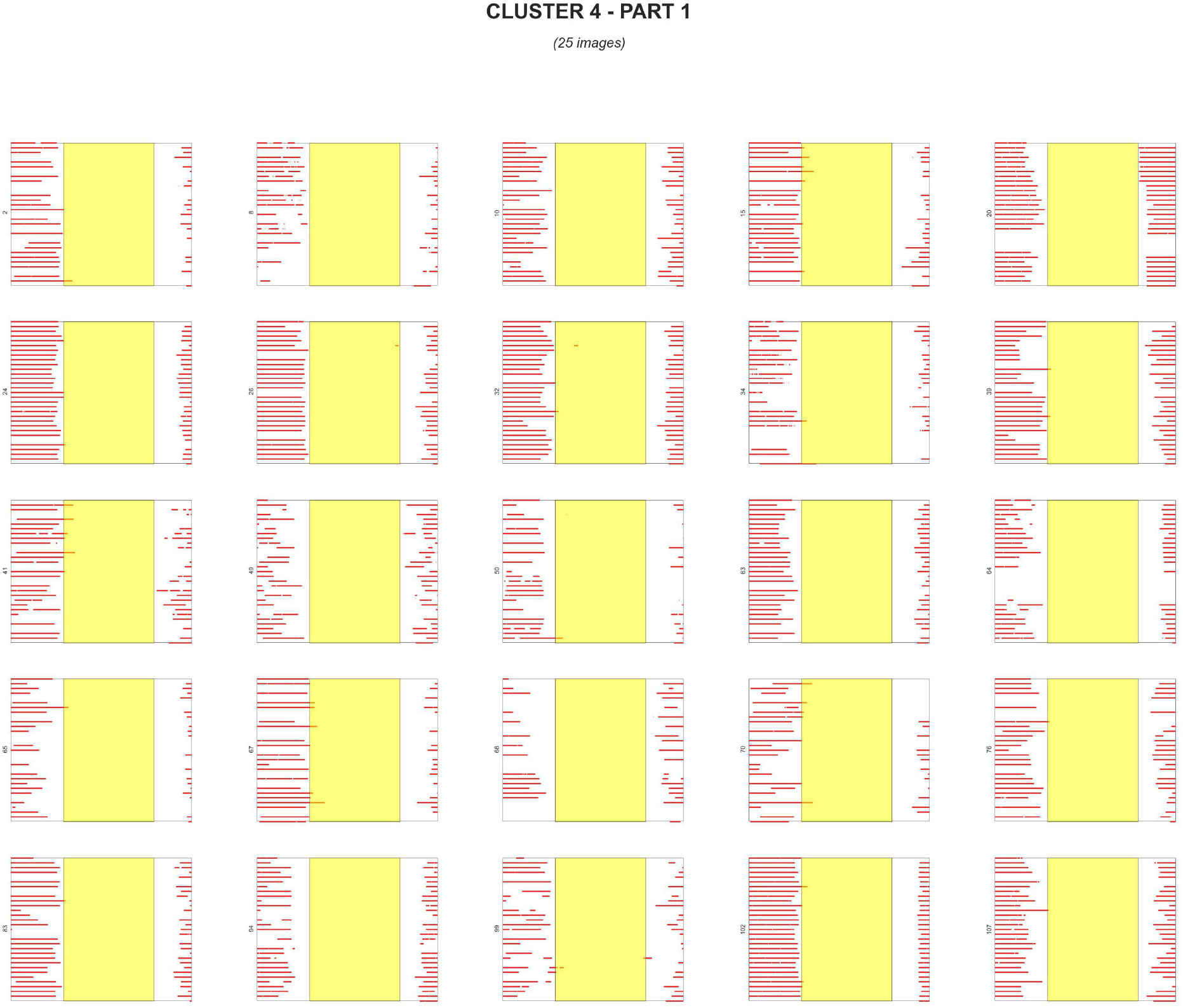
Consistent users with standard timing (n=51) - Part 1. The largest cluster shows highly consistent usage with conventional sleep timing.

**Cluster 4:**
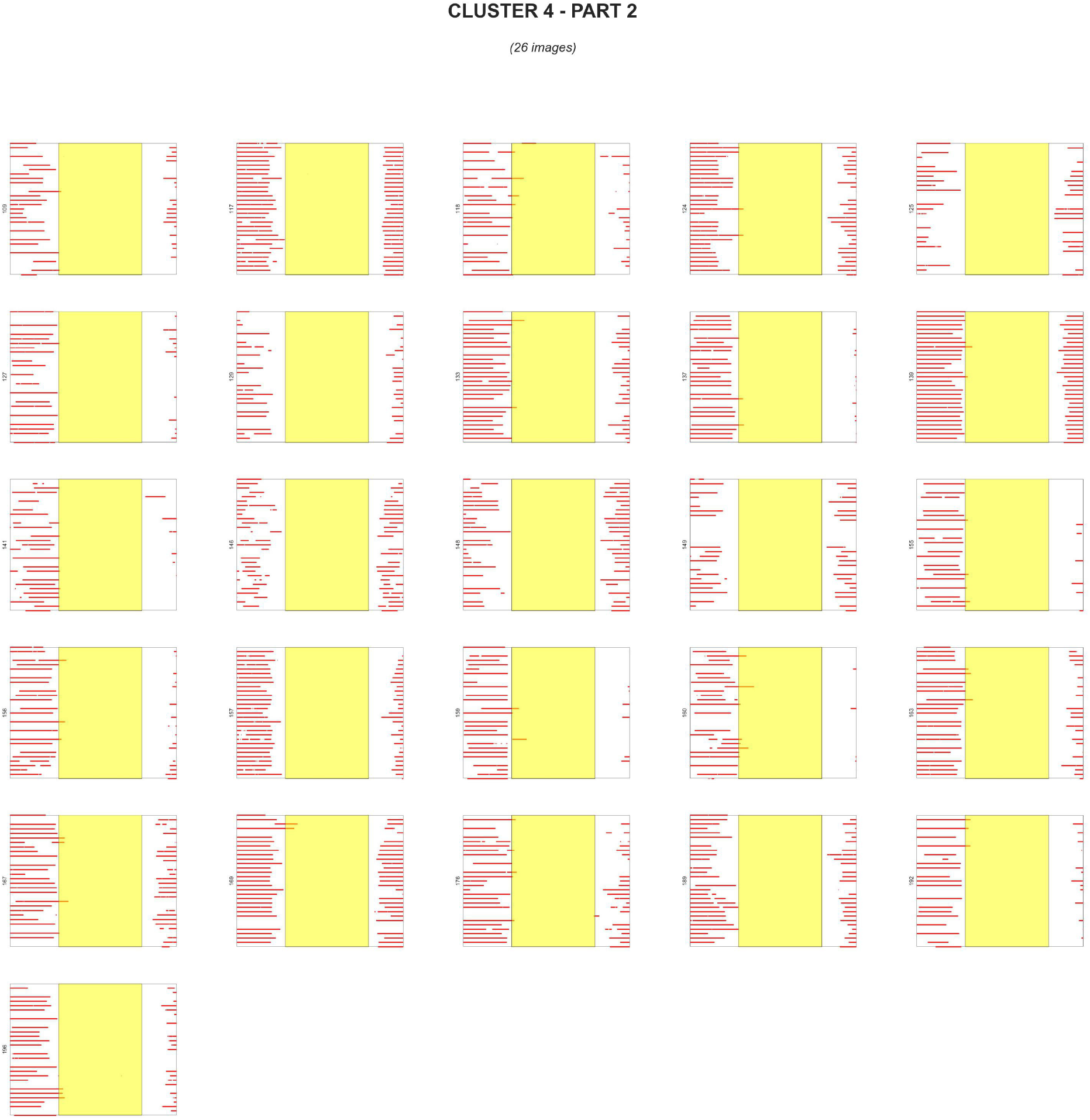
Consistent users with standard timing (n=51) - Part 2. Continuation of Cluster 4 membership.

**Cluster 5:**
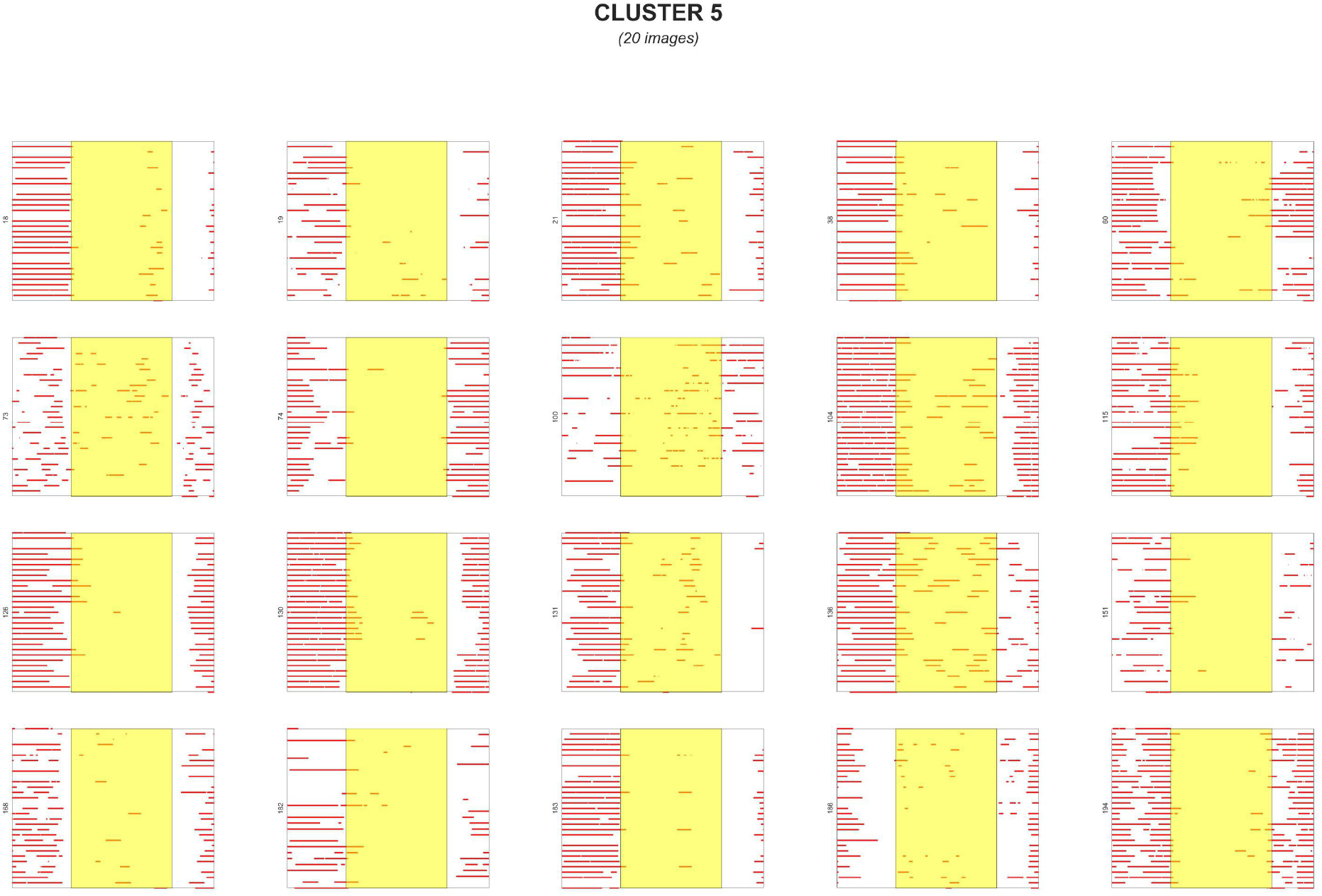
Fragmented sleep patterns (n=20). These patients show multiple on/off cycles within nights, suggesting sleep fragmentation or equipment issues.

**Cluster 6:**
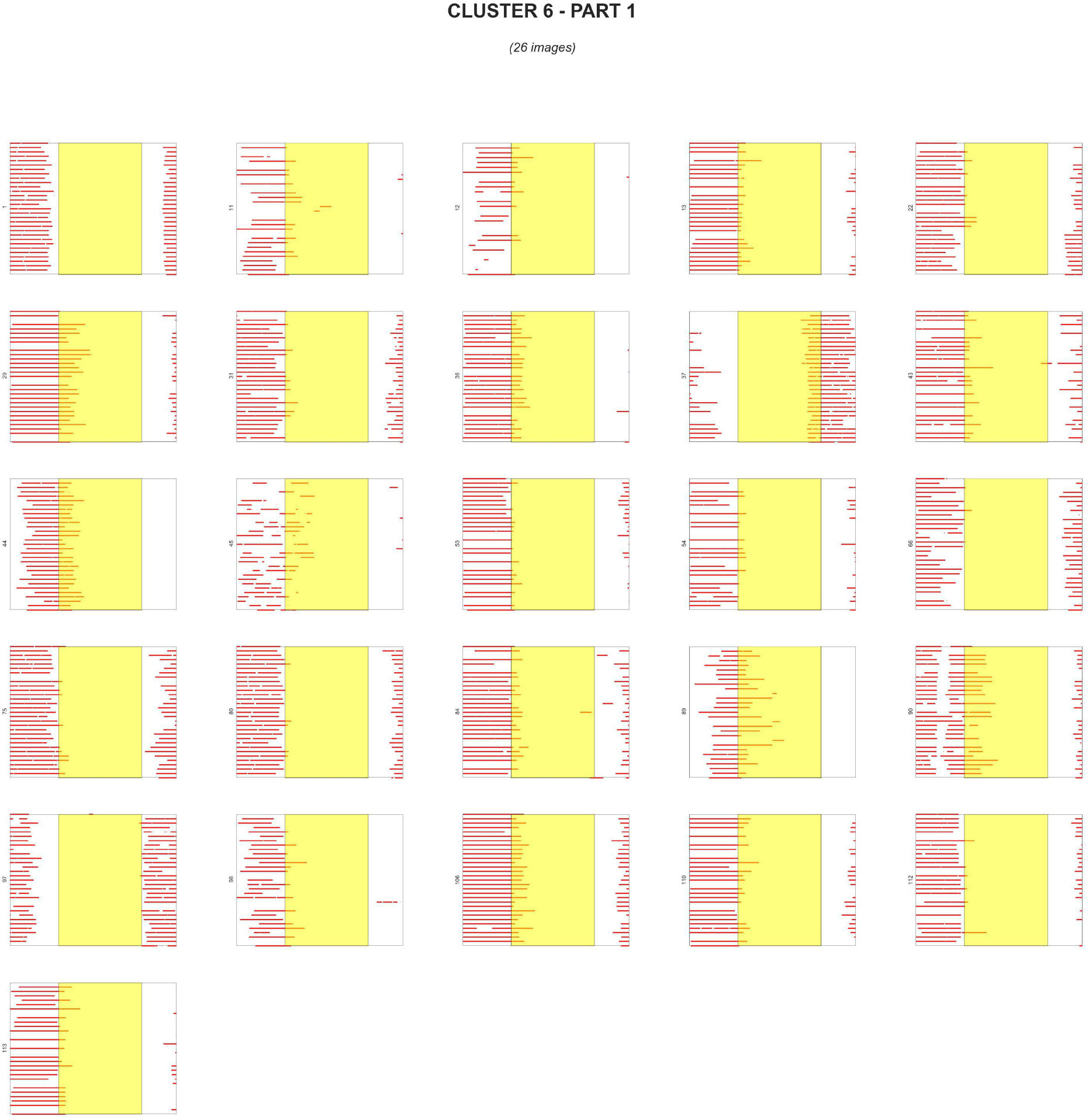
Night owls with variable adherence (n=52) - Part 1. Patients exhibit consistently late sleep onset with varying adherence quality.

**Cluster 6:**
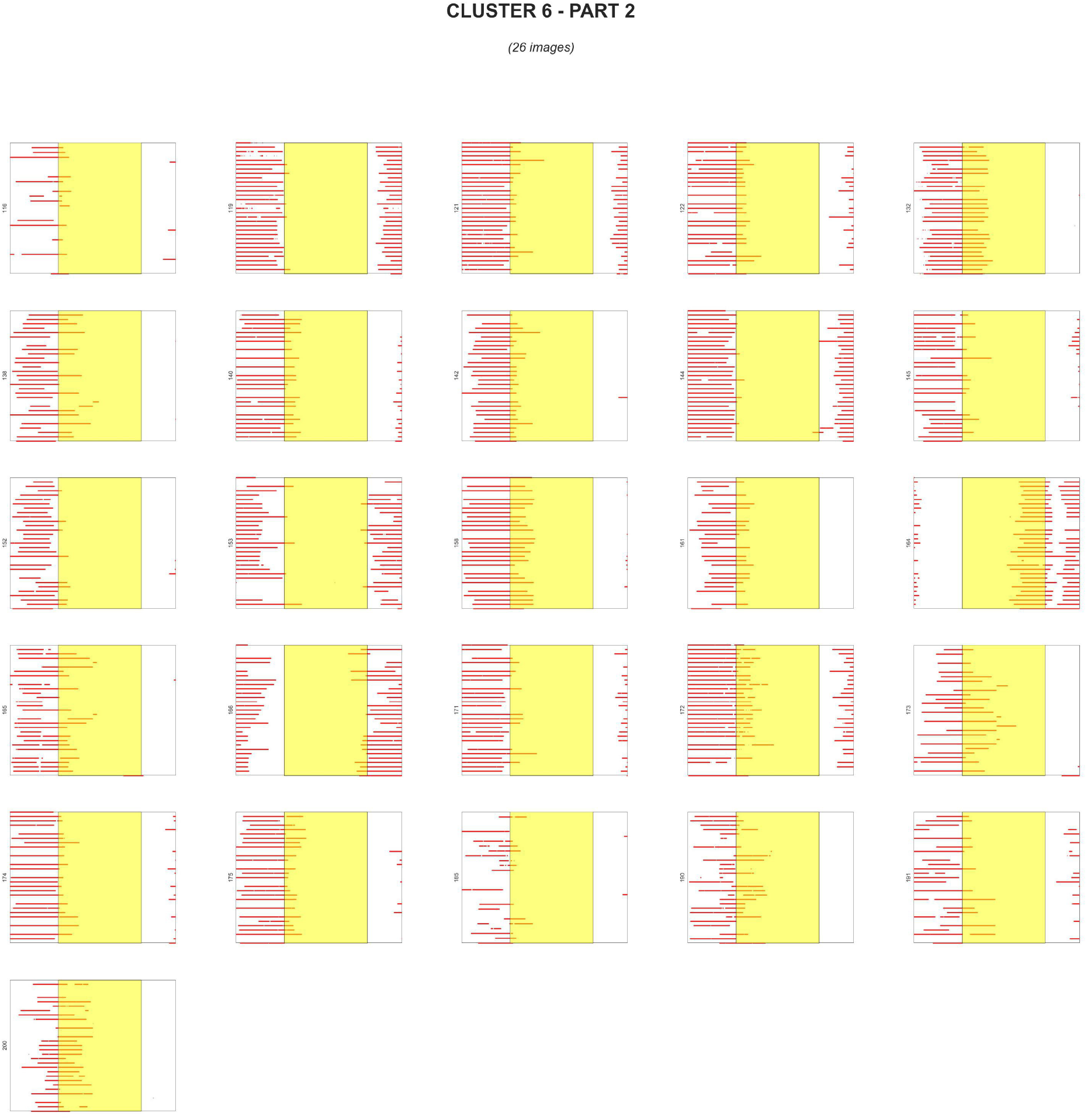
Night owls with variable adherence (n=52) - Part 2. Continuation of Cluster 6 membership.

